# Spatial transcriptomics of developing wheat seed reveals radial expression patterns in endosperm and subgenome biased expression of key genes

**DOI:** 10.1101/2025.06.08.658529

**Authors:** Tori Millsteed, David Kainer, Robert Sullivan, Xiaohuan Sun, Ka Leung Li, Likai Mao, Arlie Macdonald, Robert J Henry

**Affiliations:** Queensland Alliance for Agriculture and Food Innovation (QAAFI), University of Queensland, St Lucia 4072 QLD, Australia; ARC Centre of Excellence for Plant Success in Nature and Agriculture, University of Queensland, St Lucia 4072 QLD, Australia; Queensland Brain Institute, University of Queensland, St Lucia 4072 QLD, Australia; MGI Australia, Herston QLD 4006, Australia

**Keywords:** STOmics, seed, pericarp, embryo, marker genes, paralogs

## Abstract

Gene expression of developing seeds drives essential processes such as nutrient storage, stress tolerance and germination. However, the spatial organisation of gene expression within the complex structure of the seed remains largely unexplored. Here we report the use of the STOmics spatial transcriptomics platform to visualise spatial expression patterns in the wheat (*Triticum aestivum*) seed at the critical period of grain filling in mid seed development. We analysed >4,000,000 spatially resolved transcripts, achieving subcellular resolution of transcript localization across multiple tissue domains, and identified gene expression clusters linked to eight functional cellular groups. Notably, our analysis characterised four distinct clusters within the endosperm, which exhibited radial expression patterns from the inner to outer regions of the grain, and identified novel marker gene candidates for the clusters found. We further investigated known tissue-specific genes and identified subgenome biased expression between paralogs of puroindoline-B, metallothionein protein, and α-amylase/subtilisin inhibitor. These findings provide new detail about gene expression across and within different functional cellular groups of the developing seed and demonstrate that spatial transcriptomics could further our understanding of subgenome differences in polyploid plants.

## Introduction

The development of seeds is driven by spatiotemporally distinct gene expression, which impacts seed size and vigour of the resulting plant (Chaudhury *et al*., 2001; Ohto *et al*., 2007; Li *et al*., 2014; Bizouerne *et al*., 2021; Pelletier *et al*., 2024). Key traits such as nutrient storage, stress tolerance, germination, and the nutritional properties of seeds are influenced by gene expression in specific tissues and cellular groups, and their interactions (Dwivedi *et al*., 2021). Therefore, understanding gene expression in the complex structure of the seed is significant to plant breeders, geneticists and ecologists in the context of optimisation for food security and climate change (Dwivedi *et al*., 2021; Theissinger *et al*., 2023). While transcriptomics analyses have helped identify key genes involved in these processes, they have lacked the spatial detail needed to complete the picture of tissue-specific gene expression driving development. The emergence of spatial transcriptomics technologies has allowed for gene expression data to be placed into detailed spatial resolution, providing new insights about the genes underpinning agronomically and ecologically important traits (Yin *et al*., 2023).

Seed development is a highly coordinated process that takes place in different stages across various tissues (Bechtel *et al*., 2009). Development begins with double fertilisation resulting in formation of the endosperm and early differentiation of the embryo. For monocots the endosperm is the predominant structure of the seed and the source of energy reserves used by the embryo at germination (Brown & Lemmon, 2007). For cereal crops such as wheat (*Triticum aestivum*), endosperm development is also known as the grain filling stage and is the main determinant of yield (Sabelli *et al*., 2009). Due to its agronomic importance, grain filling has been extensively studied and it is estimated to peak between 14 and 20 days post anthesis (DPA). Endosperm development involves sequential stages of cell division and differentiation, and accumulation of storage reserves such as starch and proteins (Shewry *et al*., 2012). The process of accumulation occurs in layers with a higher protein content directed to the sub-aleurone and increasing starch deposition in the central endosperm (Shewry *et al*., 2020). Additionally, sugars and amino acids supplied via the crease are transported radially outwards towards the sub-aleurone layer and exhibit a decreasing gradient away from the crease (Ugalde & Jenner, 1990). Throughout development, wheat seeds are also photosynthetically active via the chlorophyll-rich tube- and cross-cell layers of the pericarp. However, their lack of stomata has led to the proposal that seeds utilise already respired carbon accumulating in the endosperm for photosynthesis, through a multicellular mechanism linking the endosperm and the tube- and cross-cells (Rangan *et al*., 2024). This biochemistry would be significant to yield and stress tolerance however is not yet characterised at the gene network level. Despite the established knowledge of the physiological development patterns of the wheat seed, there is a need for more detailed exploration of the spatiotemporal gene expression underlying these processes.

The polyploid status of wheat, a hexaploid plant, adds further complexity to the study of the gene expression. Modern hexaploid bread wheat resulted from hybridisation between tetraploid wheat, *Triticum turgidum* (AABB subgenomes), and diploid *Aegilops tauschii* (DD subgenome) approximately 10,000 years ago (Peng *et al*., 2011). Polyploid plants tend to have distinct traits such as larger seeds or improved vigour compared to their ancestral lines (Chan *et al*., 2022). Additionally, species with higher ploidy levels are reported to exhibit enhanced tolerance to biotic and abiotic stressors (Li *et al*., 2024). Polyploidization events often result in asymmetrical expression between subgenomes and unbalanced contribution to particular biological processes (Feldman *et al*., 2012). For example, in wheat the D subgenome is reported to be more highly expressed in response to stressors (Zheng *et al*., 2022; Powell *et al*., 2017), which has consequences for seed development. Therefore, understanding subgenome expression in wheat is agronomically important, yet very little research has considered spatial subgenome expression biases. As different tissues and cellular groups influence different developmental processes, investigating their spatial context would offer new insight about how subgenomes contribute to seed development in polyploid plants.

In the post-genomics era of genetics research, transcriptomics provides a more detailed and more cost-effective method to study gene activity in plants, uncovering countless genes influencing important traits. However, the common techniques of bulk RNA-seq and single cell RNA sequencing (scRNA-seq) lack the ability to analyse gene expression of single cells in their entire spatial context. The recent evolution of spatial transcriptomics addresses this gap (Giacomello, 2021). A number of spatial transcriptomics platforms have emerged with differing techniques to visualise gene expression, and at different resolutions. The technique of *in situ* capture and scRNA-seq are common of several platforms, allowing the entire mRNA contents of a tissue section to be sequenced with spatial information, and mapped back into the original tissue image (Yin *et al*., 2023). In recent years, this technique has been used in the study of *Arabidopsis thaliana* leaves (Xi et al, 2022), the barley grain (Pierats-llobet *et al*., 2023), the maize kernel (Fu *et al*., 2023) and ear (Wang *et al*., 2024), soybean nodules (Liu *et al*., 2023), tomato calli (Song *et al*., 2023), wheat inflorescences (Long *et al*., unpublished), and recently in the developing wheat grain (Li *et al*., 2025). Li *et al* (2025) analysed the spatial transcriptome of the developing wheat grain at 4-, 8- and 12-days post anthesis (DPA), and identified 10 distinct cell types and associated marker genes. The seed sections were cut longitudinally through the ventral and dorsal sides, providing a viewpoint of spatial expression parallel to the midline of the seed structure. Their study provided the first comprehensive spatiotemporal dataset of early wheat seed development and forms the foundation for further work to be undertaken at different time points and with new genetic targets.

Here we utilised STOmics scStereo-seq to analyse the spatial transcriptome of the 14 DPA wheat seed. Our study provides the continuation of the developmental time points highlighted by Li *et al* (2025), focusing on the peak of grain filling in mid seed development. We also investigated the alternative view of spatial gene expression with tissue sections cut longitudinally through the lateral sides of the seed, resulting in cross sections parallel with the dorsal and ventral sides of the seed structure. Our study aimed to assess spatial gene expression patterns in the complex structure of the seed during grain filling, as well as subgenome biased expression of key genes.

## Results

STOmics scStereo-seq was used to measure spatial gene expression in the developing wheat seed. Sections of 14 DPA seeds were cut in a cryostat to a thickness of 20 µm and mounted onto the STOmics transcriptomics assessment chips, covered in an array of spatially tagged DNA nanoballs (DNBs). The contents of the chips were sequenced, aligned to the reference genome IWGSC CS RefSeq v2.1 (Zhu *et al*., 2021), and mapped back into their original locations. The resulting spatially tagged gene expression matrix was also overlaid with fluorescent microscope images of the original tissue, to place the spatial expression patterns within the cell and tissue architecture. Due to challenges adhering the tissue sections to the chip surface, the most intact section was chosen as the focus for this study, though some folding of the tissue is present. Data from replicate chips have been used to support our findings where possible.

### Gene expression matrix reveals areas of high gene activity that align with key tissue areas

Fluorescent microscope images of the seed sections highlighted the tissue structure. Well-defined cell walls of the pericarp and aleurone layers were visible in the blue channel, and a scattered array of cell nuclei of varying sizes in the endosperm, aleurone and pericarp, were visible in the green channel (Figure 1AB). The gene expression matrix was visualised as a heat map which depicted areas of low expression in dark blue and areas of high expression in red. The areas of highest gene expression were in the embryo, the crease, the inner endosperm, and the pericarp (Figure 1CD). Additionally, gene expression levels were not symmetrical on both sides of the seed, with both the pericarp and endosperm on one side (lower in our figure) exhibiting greater expression levels than the other side.

**Figure 1:**
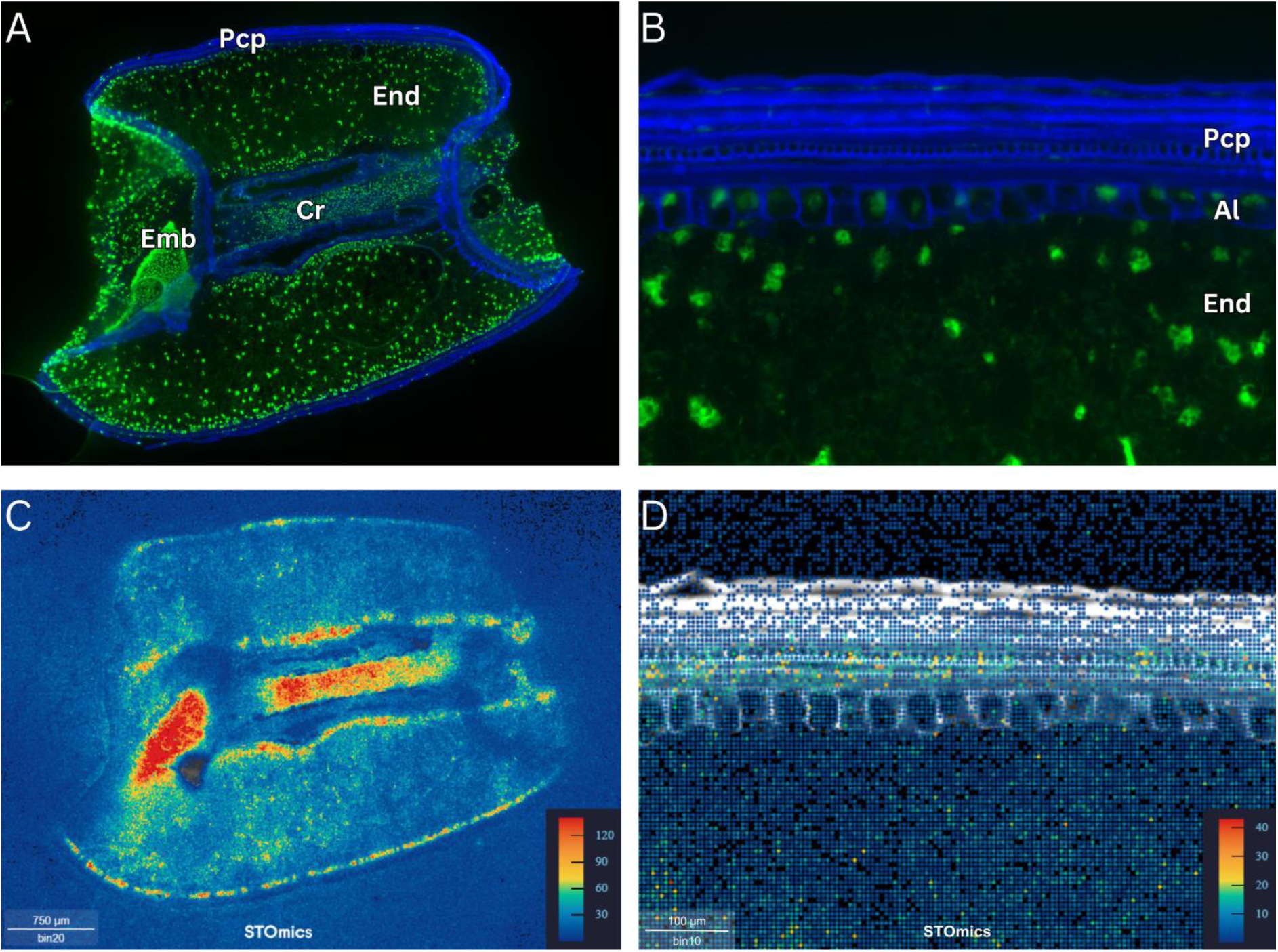
Fluorescent microscope images and gene expression heat maps of 14 DPA wheat seed section. **A)** Image of wheat seed section, with cell walls in blue and cell nuclei in green, showing pericarp (Pcp), endosperm (End), embryo (Emb) and crease (Cr) tissues. **B)** Image showing enhanced detail of pericarp, aleurone (Al) and endosperm tissues. **C)** Gene expression heat map of the same section at bin20 resolution, showing highest gene expression in red and lowest gene expression in blue (MID count per bin). **D)** Corresponding gene expression heat map of magnified image B reveals the specific spatial distribution of the anatomically well-defined layers of the pericarp, aleurone and endosperm layers at bin10 resolution. Scale bars in A, C = 750 µm and B, D = 100 µm.

For the STOmics chip containing the most intact tissue section there were 92,046,597 DNBs located beneath the tissue area. Of these, 3,273,120 DNBs successfully captured 4,244,343 MIDs (unique molecular identifiers) at subcellular resolution, resulting in an average of 1.3 MIDs from an average of 1.09 genes per DNB (Figure S2). The sparseness of the capture array was alleviated by grouping adjacent DNBs into 20x20 bins (Bin20) where each Bin20 contained an average of 18.47 MIDs from 14 genes. For the replicates analysed there were between 3,237,120 and 11,599,308 DNBs that successfully captured mRNA beneath the tissue areas, resulting in an average capture array of 1.30 – 1.57 MIDs per DNB (Figures S2, S4, S6).

### Clusters identified match key tissue areas and show radial expression patterns

Spatial Leiden clustering was performed on the expression data from each chip at a resolution of Bin50. A total of 8 clusters were identified and linked to functional cellular groups (Figure 2). These were the pericarp (Figure 2B), the sub-aleurone (Figure 2C), the central endosperm (Figure 2D), the inner endosperm boundary (Figure 2E), the inner endosperm ring (likely transfer cells) (Figure 2F), the crease (Figure 2G), the inner crease (Figure 2H) and the embryo (Figure 2I). While clusters B, G, H and I were linked to a single tissue type or cellular group, clusters C, D, E and F were all located within the same tissue area, the endosperm. The sub-aleurone cluster formed the outermost layer of the endosperm and followed the ring-like shape of the outer boundary of the tissue section, with some expression overlapping with the central endosperm. The central endosperm cluster covered the largest area of the seed section and, while strongly localised to the central endosperm, also exhibited some overlap with the neighbouring clusters. The inner endosperm ring and inner endosperm boundary clusters also appeared in a radial shape, encircling the crease at the centre of the tissue section. The inner endosperm ring cluster was one of the areas of greatest gene expression in the seed, and the presence of the inner endosperm boundary cluster, distinct from the inner endosperm ring, highlighted an area of unique gene expression in the endosperm that hasn’t previously been characterised. Similar clusters exhibiting radial expression patterns, and distinct expression of the inner endosperm ring, were also identified in the replicate chips (Figure S8).

**Figure 2:**
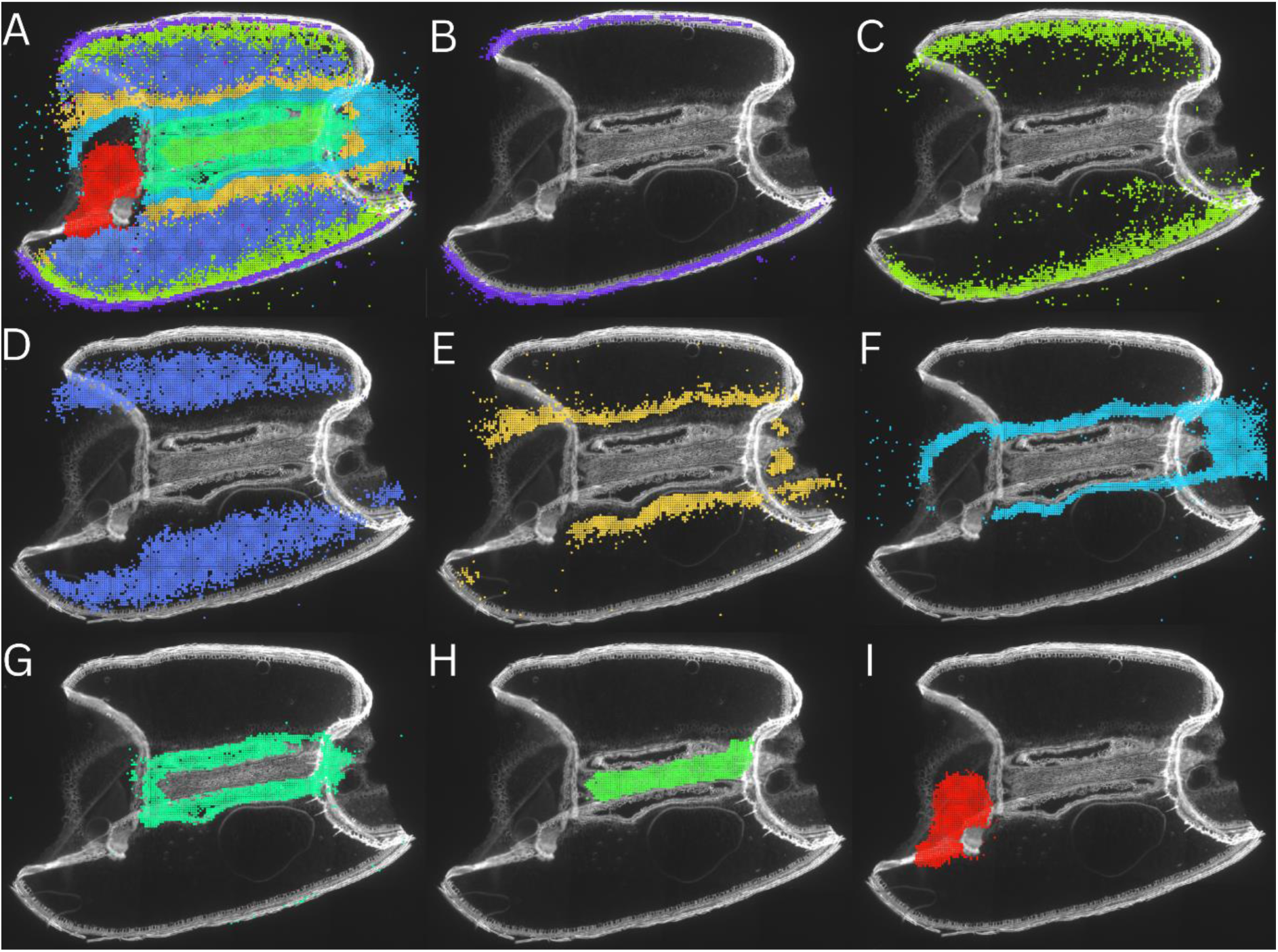
Gene expression clusters identified by spatial Leiden clustering in 14 DPA wheat seed section. **A)** Visualisation of all major gene clusters, **B)** pericarp cluster, **C)** sub-aleurone cluster, **D)** central endosperm cluster, **E)** inner endosperm boundary cluster, **F)** inner endosperm ring cluster, **G)** crease cluster **H)** inner crease cluster and **I)** embryo cluster.

### Expression of known tissue-specific genes confirms data accuracy

We analysed the spatial expression of known tissue-specific genes to validate the robustness of our data and confirm their expression in the expected tissue types. This included genes known, or expected, to be expressed in the pericarp, endosperm and embryo tissues. The genes for puroindoline-B were investigated, as they are related to wheat endosperm hardness and known to be highly expressed in the endosperm and aleurone cells (Nirmal *et al*., 2016). This was confirmed by our gene expression matrix which showed high levels of expression in the endosperm compared to other tissue types, particularly the crease (adjusted approximate permutation tests p < 0.01). However, our data also showed that these genes were expressed at some level in all tissue types (Figure 3ABC). *TaNAC019*, a wheat transcription factor involved in starch biosynthesis in the endosperm (Gao *et al*., 2021) was also investigated, and expression levels were found to be generally low. These genes appeared to be mostly localised to the endosperm (Figure S9), however, this wasn’t found to be statistically significant. Similarly, wheat transcription factor *TabZIP28,* also known to be involved in starch biosynthesis, (Song *et al*., 2020), appeared to be mostly expressed in the endosperm (Figure S10), but expression levels were too low for this difference to be considered significant.

**Figure 3.**
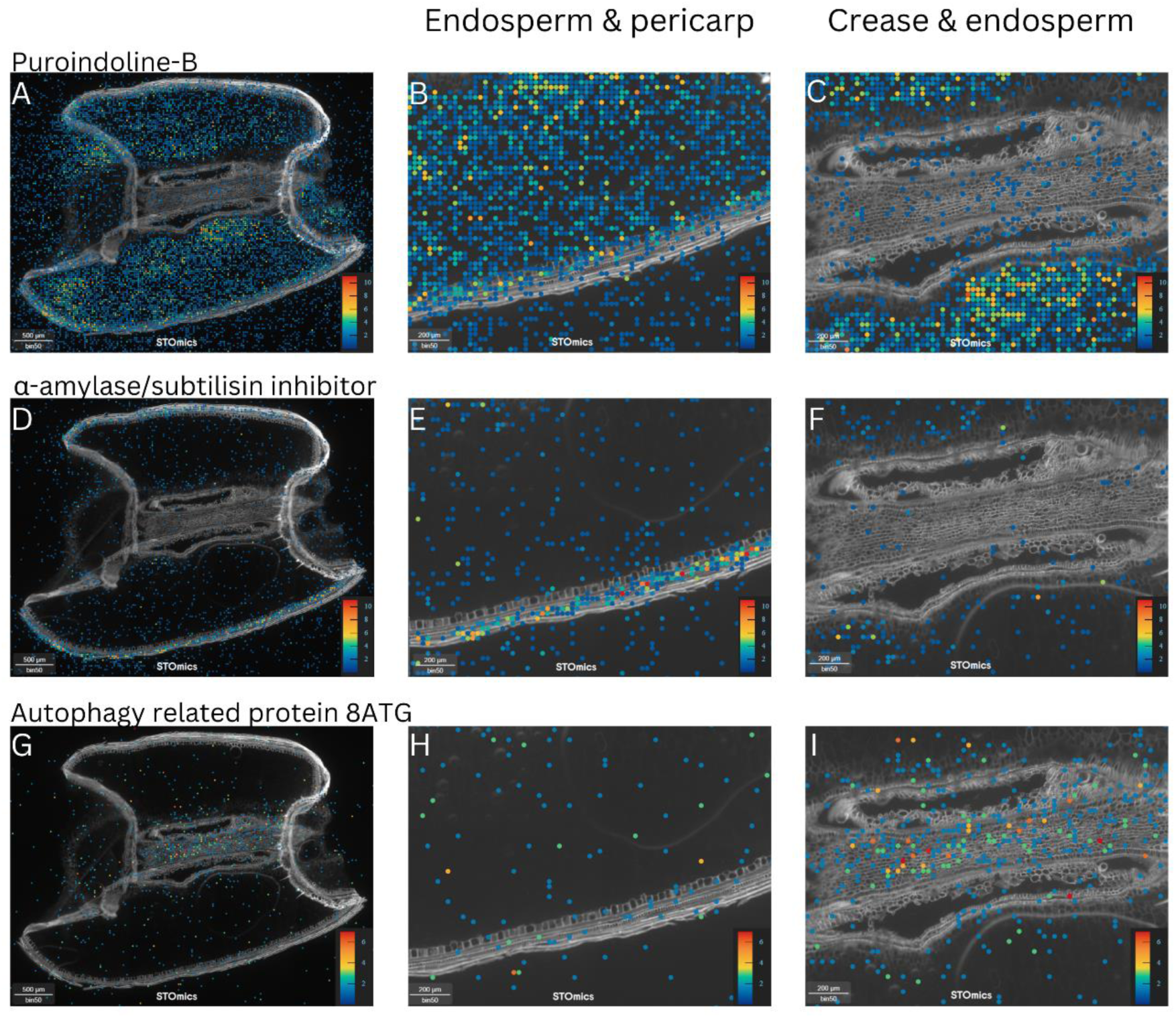
Expression of known tissue-specific genes in various tissues of the 14 DPA wheat seed section. **A)** Puroindoline-B expression in entire grain section, **B)** Puroindoline-B expression in endosperm and pericarp, **C)** Puroindoline-B expression in crease and endosperm, **D)** α-amylase/subtilisin inhibitor expression in entire grain section, **E)** α-amylase/subtilisin inhibitor expression in endosperm and pericarp, **F)** α-amylase/subtilisin inhibitor expression in crease and endosperm, **G)** Autophagy related protein 8ATG expression in entire grain section, **H)** Autophagy related protein 8ATG expression in endosperm and pericarp, **I)** Autophagy related protein 8ATG expression in crease and endosperm.

Metallothionein protein genes were assessed as these were expected to be embryo specific (Kawashima *et al*., 1992). These were most highly expressed in the embryo, followed by the pericarp and crease, and then lower expression in the endosperm tissue (all adjusted approximate permutation tests p < 0.01) (Figure S11). The genes encoding the EM promoter protein were also investigated as they are known to be specifically expressed in the embryo and the aleurone layer (Furtado & Henry, 2005). These genes were most highly expressed in the embryo and the pericarp, and they also exhibited some lower expression in the endosperm and crease (all adjusted approximate permutation tests p < 0.01) (Figure S12).

Genes encoding α-amylase/subtilisin inhibitor were investigated, as they are highly expressed in the barley pericarp (Furtado *et al*., 2003). These genes exhibited a pattern of strongest expression in the pericarp in wheat (all adjusted approximate permutation tests p < 0.01), as well as expression at lower levels throughout the endosperm, crease and embryo (Figure 3DEF). The genes for pyruvate orthophosphate dikinase were also assessed as pericarp specific, due to their role in photosynthesis (Rangan *et al*., 2016). These genes were most highly expressed in the pericarp and the embryo (all adjusted approximate permutation tests p < 0.01), but also showed some dispersal in the endosperm (Figure S13). Lastly, the genes encoding autophagy related protein 8ATG were investigated. These genes are highlighted in the literature as being involved in programmed degradation of pericarp cells (Li *et al*., 2021). While there was some expression of these genes in the pericarp layer, expression appeared to be more strongly localised to the crease region of the grain, with some expression in the endosperm at similar levels to the expression seen in the pericarp (Figure 3GHI) (Table 1). However, these observed differences were not found to be statistically significant.

**Table 1.** Known tissue-specific genes expected and observed spatial expression patterns, and subgenome specific expression in 14 DPA wheat seed section. Includes gene name and/or function, NCBI Gene ID, subgenome, expected spatial expression based on the literature, spatial expression observed in the data, and the overall paralog expression (MID count).

| Gene name/<br>function | Gene ID | Sub-<br>genome | Expected<br>spatial<br>expression | Observed<br>spatial<br>expression<br>in our data | Overall paralog<br>expression (MID count) |  |  |
| --- | --- | --- | --- | --- | --- | --- | --- |
|  |  |  |  |  | Rep 1 | Rep 2 | Rep 3 |
| Puroindoline-B | LOC123101925 | A | Endosperm/<br>aleurone | Endosperm/<br>aleurone | 2836 | 15,021 | 3817 |
|  | LOC543308 | B |  |  | 7387 | 35,537 | 10,749 |
|  | LOC100125699 | D |  |  | 24,634 | 97,257 | 34,577 |
| TaNAC019 | LOC123057832 | A | Endosperm | Endosperm | 238 | 373 | 226 |
|  | LOC123064838 | B |  |  | 0 | 0 | 0 |
|  | LOC123073994 | D |  |  | 82 | 262 | 86 |
| TabZIP28 | LOC123187748 | A | Endosperm | Endosperm/<br>crease | 153 | 219 | 146 |
|  | LOC123043995 | B |  |  | 287 | 383 | 200 |
|  | LOC123051864 | D |  |  | 205 | 242 | 148 |
| Metallothionein-protein |  | A | Embryo | Embryo/<br>pericarp/<br>crease | 0 | 0 | 0 |
|  | LOC123104584 | B |  |  | 2131 | 585 | 785 |
|  | LOC123179983 | D |  |  | 21,397 | 8362 | 9896 |
| EM promoter<br>protein | LOC543476 | A | Embryo | Embryo/<br>pericarp/<br>crease | 1543 | 1617 | 893 |
|  | LOC543084 | B |  |  | 2100 | 1292 | 819 |
|  | LOC123182837 | D |  |  | 1439 | 894 | 550 |
| α-amylase/<br>subtilisin<br>inhibitor | LOC123185730 | A | Pericarp | Endosperm | 123 | 458 | 109 |
|  | LOC123041664 | B |  | Pericarp/ | 4286 | 13,229 | 3443 |
|  | LOC123049628 | D |  | endosperm | 2730 | 13,649 | 4056 |
| Pyruvate<br>orthophosphate<br>dikinase | LOC123055206 | A | Pericarp | Pericarp/ | 1380 | 3111 | 1038 |
|  | LOC123132411 | B |  | embryo/ | 609 | 1658 | 579 |
|  | LOC123181969 | D |  | endosperm | 1418 | 2625 | 884 |
| Autophagy<br>related protein<br>8ATG | LOC123188447 | A | Pericarp | Crease/ | 533 | 958 | 371 |
|  | LOC123044698 | B |  | endosperm/ | 711 | 1021 | 432 |
|  | LOC542962 | D |  | pericarp | 690 | 1123 | 515 |

### Subgenome biased expression identfied for some key genes

Subgenome biased expression was identified for some of the genes analysed. For puroindoline-B the subgenome D paralog was significantly more highly expressed overall than the subgenome A and B paralogs (Figure 4ABC), with 24,634 MIDs compared to 7387 and 2836 respectively (all adjusted approximate permutation tests p < 0.01). The A and B paralogs were also found to be significantly differently expressed (all adjusted approximate permutation tests p < 0.01). Similarly, for metallothionein protein, the subgenome D paralog was expressed at significantly higher levels than the subgenome B paralog overall (Figure 4GH), with 21,397 MIDs compared to 2131 MIDs (all adjusted approximate permutation tests p < 0.01). There was also notably no subgenome A paralog of this metallothionein protein isoform present in the reference genome (Zhu *et al*., 2021).

**Figure 4.**
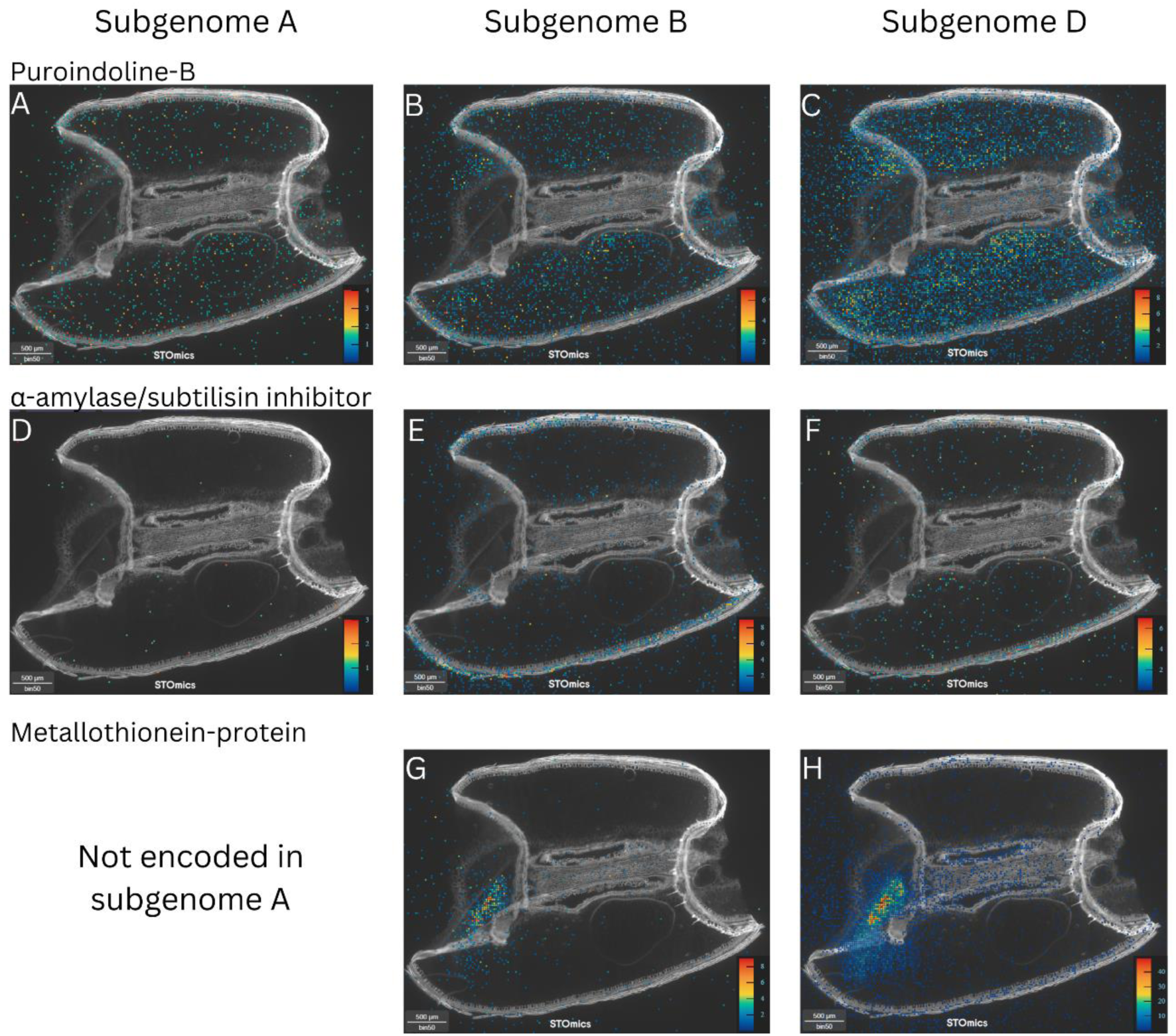
Subgenome biased expression levels of paralogs of known tissue-specific genes in 14 DPA wheat seed section. **A)** Puroindoline-B subgenome A expression, **B)** Puroindoline-B subgenome B expression, **C)** Puroindoline-B subgenome D expression, **D)** α-amylase/subtilisin inhibitor subgenome A expression, **E)** α-amylase/subtilisin inhibitor subgenome B expression, **F)** α-amylase/subtilisin inhibitor subgenome D expression, **G)** Metallothionein-protein subgenome B expression, **H)** Metallothionein-protein subgenome D expression.

The genes for α-amylase/subtilisin inhibitor exhibited subgenome biased spatial expression (Figure 4DEF). The subgenome A paralog was expressed at significantly lower levels overall than the subgenome B and D copies with 123 MIDs compared to 4286 and 2730 MIDs respectively (all adjusted approximate permutation tests p < 0.01). The subgenome A paralog also appeared to be nearly only expressed in the endosperm, though this observed difference was not found to be statistically significant. Conversely, the subgenome B and D paralogs were significantly more highly expressed in the pericarp tissue than any other tissue type (all adjusted approximate permutation tests p < 0.01).

### Novel marker genes identified for gene expression clusters

The five strongest marker gene candidates for each cluster were identified (Figure 5). Strong marker genes were identified for the pericarp, the crease, the embryo and the inner endosperm ring, while a number of marker genes were identified for the endosperm but were not specific to a particular endosperm cluster. For the pericarp, the strongest marker gene candidates were LOC123149767 (26 kDa endochitinase 2) and LOC123108591 (non-specific lipid-transfer protein 2P-like), which were nearly exclusively expressed in the pericarp (Figure 5), and were also identified as marker genes for this tissue in the replicate chips (Figures S20, S21). The strongest marker gene candidate for the crease and inner crease clusters was LOC123104148 (induced stolen tip protein (TUB8)) which was also supported by the replicates (Figures S20, S21). For the embryo cluster, the strongest marker gene was LOC123113644 (cupincin-like) and was also identified as a marker for this tissue in the replicate chips (Figure S21).

**Figure 5.**
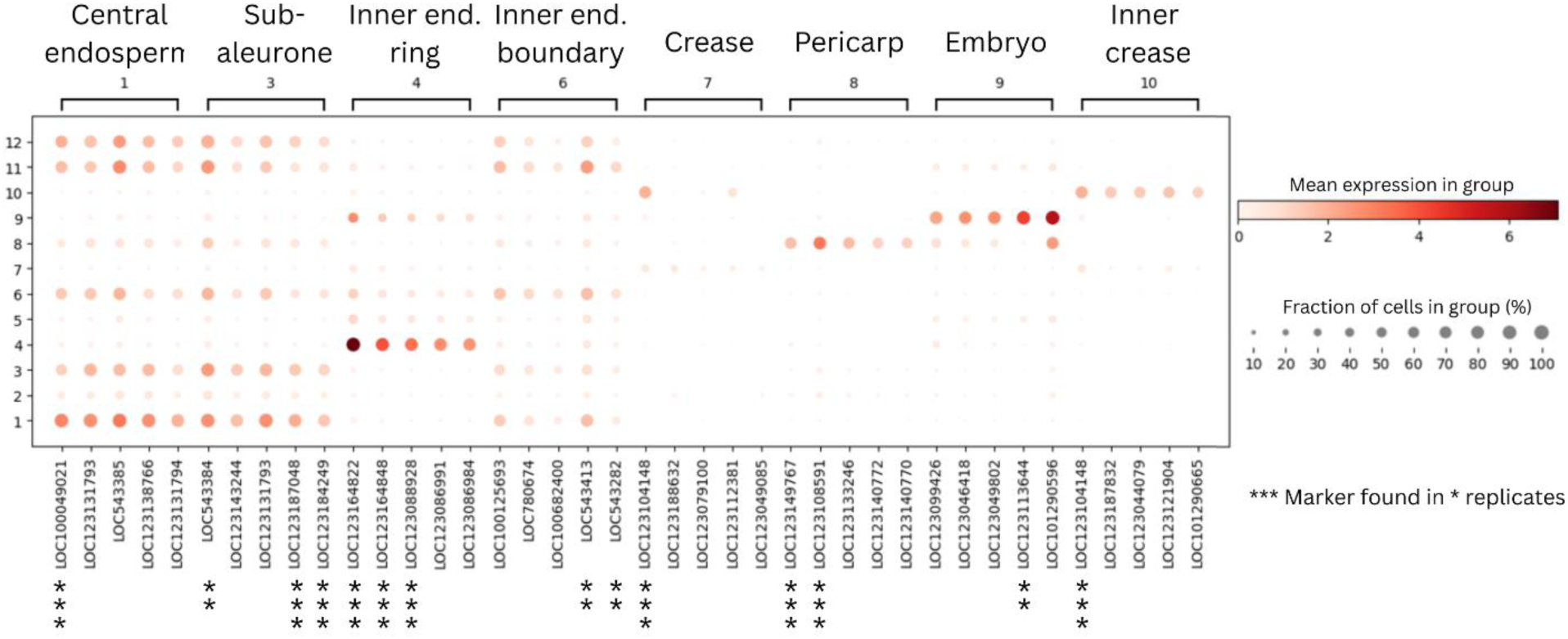
Marker gene plot of the five strongest marker gene candidates for each gene expression cluster, and alignments depicting the expression of each gene in other clusters. Dot size represents the percentage of cells (bins) in the cluster the gene is found in, and dot colour represents the mean expression (MID count) of the gene in those cells. from the Astrix (*) represent the number of times a gene was identified as a marker for its tissue type across three replicate chips. The columns for clusters 2, 5, 11 and 12 have been removed as they were determined to be background noise, or a result of tissue folding and were not linked to any functional cellular group.

The strongest marker genes for the inner endosperm ring cluster were LOC123164822, LOC123164848 and LOC123088928, which were identified in all replicates (Figures 5, S20, S21). The gene LOC123164822 was the most highly expressed in the gene expression matrix overall, and the three were highly similar (94.75-97.95% nucleotide similarity) (Sayers *et al*., 2025), however, all three are ‘uncharacterised’ in the reference genome (Zhu *et al*., 2021). These genes were predicted to encode Bifunctional inhibitor/plant lipid transfer proteins/seed storage helical domain-containing proteins, by automatic annotation (Bateman *et al*., 2024). The most similar gene to LOC123164822 that is characterised in the reference genome, with approximately 94% similarity, is known as Endosperm transfer cell specific PR60 precursor (Zhu *et al*, 2021). Additionally, Blastn analysis found a similar gene in the Emmer wheat genome known as putative lipid transfer protein precursor (PR60) gene (accession number FJ459807.1), and BLASTx analysis revealed a structurally similar protein in the *Arabidopsis thaliana* genome known as Bifunctional inhibitor/lipid-transfer protein/seed storage 2S albumin superfamily protein (accession number 834904), with approximately 90% similarity (Sayers *et al*., 2025). Therefore, we have inferred that these marker genes are most likely lipid transfer protein genes.

All marker gene candidates identified for the central endosperm, sub-aleurone and inner endosperm boundary clusters were specific to the endosperm but were expressed at similar levels across neighbouring endosperm clusters (Figure 5). These included genes for α-, β- and γ-gliadins and α-amylase/trypsin inhibitors (Table 2).

**Table 2.**
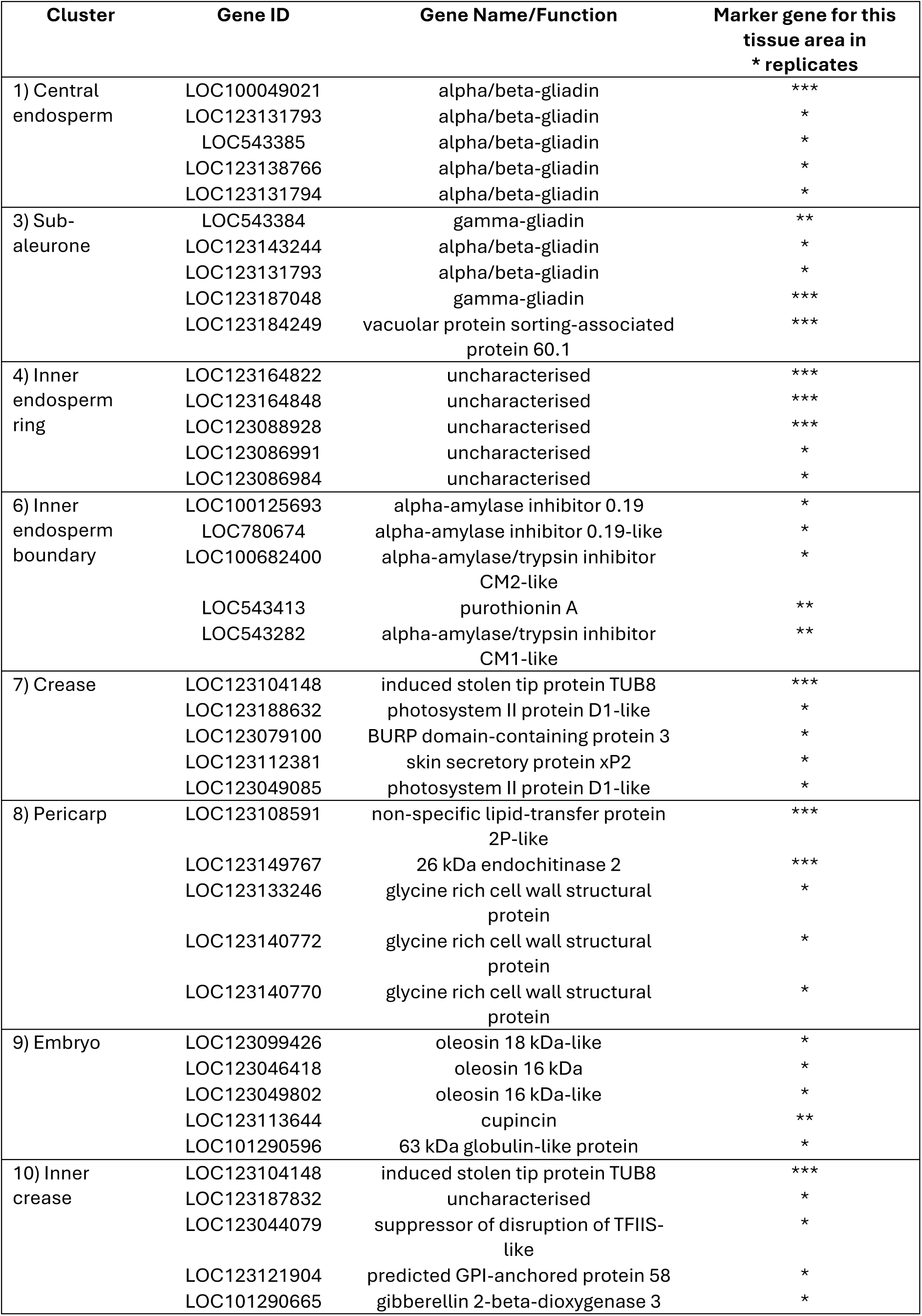
Marker genes identified for each gene expression cluster, showing the cluster name, NCBI Gene ID, the gene name/function, and the number of times a marker gene was identified for a specific cluster across three replicate chips (*).

## Discussion

Transcriptomics analyses of the wheat seed have previously uncovered key genes involved in seed development, stress tolerance, pathogen resistance, baking quality, and overall yield (Yu *et al*., 2016; Powell *et al*., 2017; Rangan *et al*., 2017; Henry *et al*., 2018; Rangan *et al*., 2020; Ahmadi-Ochtapeh *et al*., 2024). These studies have advanced optimization of the wheat crop as a food resource, however, have lacked the capacity to link genes to functional cellular groups in their detailed spatial context. Li *et al* (2025) published the first comprehensive spatiotemporal dataset of early seed development in wheat and identified 10 distinct cell types and associated marker genes. In the present study, spatial transcriptomics of the 14 DPA wheat seed revealed gene expression clusters linked to functional cellular groups, including the pericarp, endosperm, embryo and crease tissues, and identified novel marker genes linked to the clusters found. Our data also highlighted the utility of this technology to investigate subgenome biased spatial expression, highlighting expression differences between paralogs of key genes.

### Gene expression clusters of the endosperm exhibit radial pattern

The physiological stages of wheat seed development have been established to take place in radial layers, where protein and starch accumulation differ between the outer and inner regions of the endosperm (Shewry *et al*., 2020), and transport of nutrients from the crease follows a radial gradient outwards towards the sub-aleurone (Ugalde & Jenner, 1990). Between approximately 8-21 DPA the endosperm transitions from a liquid to a more solid state as this process of deposition takes place. The peak of grain filling is estimated to occur at 14-20 DPA, making this period significant to the yield and vigour of the resulting seed (Shewry *et al*., 2012). Li *et al*’s, (2025), analysis of the early developing wheat seed identified 10 gene expression clusters including the inner and outer pericarp, the testa, the embryo and embryo surrounding regions, the aleurone and sub-aleurone layers, the central cells of the starchy endosperm, the transfer cells surrounding endosperm, and the cavity fluid. From 4-12 DPA the pericarp layers made up the largest area of the seed sections and the endosperm was not yet cellularised, though multiple endosperm specific clusters could still be identified. In this study the main viewpoint of spatial gene expression was of a longitudinal cross section parallel to the midline of the seed structure, which revealed a ring-like shape of the aleurone and sub-aleurone clusters. However, the other endosperm clusters didn’t appear to follow a radial pattern from this view.

In the present study of the 14 DPA wheat seed, the endosperm region accounted for the largest area, highlighting the significance of the 12-14 DPA period for endosperm development and grain filling. Eight gene expression clusters were identified that largely corresponded with the clusters identified by Li *et al* (2025), including the pericarp, sub-aleurone, central endosperm, inner endosperm ring (transfer cells), and the embryo clusters. However, in the more mature grain only one pericarp cluster was identified, and the further development of the endosperm allowed for four distinct gene expression clusters to be delineated. These included the sub-aleurone, the central endosperm, and the inner endosperm ring clusters, as well as the ‘inner endosperm boundary’ cluster, which we suggest forms a boundary region between the starchy endosperm cells and the transfer cells. It should also be noted that the ‘embryo surrounding region’ cluster was discounted from our data as this area was affected by tissue folding. Our study highlighted the opposing view of spatial gene expression, with longitudinal cross-sections parallel to the dorsal and ventral sides of the seed structure. This revealed that the endosperm specific clusters formed ring-like patterns, following the oval shape of the seed and radiating inwards to encircle the crease. These gene expression patterns appear to align with the physiological development patterns of the seed, where different stages of protein, starch and nutrient deposition are taking place in layers across the endosperm at one time point. This study is the first time that four distinct regions of gene expression could be delineated from the endosperm during the peak of grain filling.

Interestingly, the gene expression matrix also revealed differences in overall expression between the different sides of the seed, both in the endosperm and pericarp tissues. We suggest that this could be a result of the position on the plant, with one side of the seed subject to more direct light resulting in higher biochemical activity and gene expression. This finding highlights the nuance of gene expression patterns that can be captured by spatial transcriptomics, as the gene expression of cells side by side can be compared and quantified in their entire spatial context.

### Spatial transcriptomics highlights subgenome biased expression patterns

In polyploid plants, the contribution of subgenomes to different developmental and regulatory traits can be unbalanced, making subgenome specific gene expression an important aspect of transcriptomics analyses. Differences in expression between subgenome copies of various genes are reported in wheat. It is thought that unbalanced expression between paralogs is the first stage of neo-functionalisation and enables functional specialisation and adaptability in polyploid plants (Ramirez-Gonzalez *et al*., 2018). For instance, the B, and D subgenomes are known to be more highly expressed in wheat overall (Ramirez-Gonzalez *et al*., 2018), and subgenome D is reported to contribute disproportionately to abiotic and biotic stress responses (Zheng *et al*., 2022; Powell *et al*., 2017). Here, we reported subgenome specific differences in expression between homologous genes triplicates of puroindoline-b, metallothionein protein and α-amylase/subtilisin inhibitor. These findings were consistent with the literature as the subgenome B and D copies were more highly expressed than the subgenome A copies, when they were present. However, there was no subgenome A copy for this metallothionein protein isoform encoded in the reference genome (Zhu *et al*., 2021), which could be an example of a gene lost due to unbalanced subgenome expression, making it vestigial.

Similarly, some subgenome specific spatial expression patterns have been reported in wheat (Wang *et al*., 2021), but are not yet well characterised as the capacity for detailed spatial transcriptomics has only recently advanced. Li *et al* (2025) investigated subgenome biased spatial expression of the B3 transcription factor family in the 8 DPA seed section. They found higher expression of the B subgenome in the endosperm and aleurone, as well as greater expression of the subgenome B copy of *TaAB13-B1* in the embryo, at this developmental stage. Here, we reported subgenome specific spatial expression patterns for α-amylase/subtilisin inhibitor, where the subgenome A copy was expressed at low levels in all tissues, compared to the subgenome B and D copies, which were significantly more highly expressed in the pericarp than any other tissue. Therefore, our study provides further evidence of differentiation between subgenomes at the spatial gene expression level. These findings are significant in the context of genetic research to optimise seed traits, as they highlight the utility of spatial transcriptomics for target gene analysis with both subgenome and tissue specificity.

### Marker genes highlight functions of key cellular groups

The functions of particular cellular groups or tissue types, and how these components interact to influence important developmental traits, can be indicated by the genes most highly expressed in those tissues. In our study, strong marker genes were identified for the pericarp, crease, embryo and inner endosperm ring clusters, meaning they were highly expressed in those tissues and comparatively little in others. Additionally, marker genes were identified for the endosperm tissue, but weren’t specific to a particular endosperm cluster. These findings have provided new information about the activity in these cellular groups during mid seed development.

The pericarp region is a structurally significant component of the seed and serves as an important protective layer for the endosperm and embryo tissues during development (Bechtel *et al*., 2009). The strongest marker genes identified in the pericarp were LOC123149767, encoding endochitinase, and LOC123108591, encoding a non-specific lipid-transfer protein. Endochitinases are known to play an important role in plant defence against fungal pathogens (Vaghela *et al*., 2022), while non-specific lipid transfer proteins have roles in cell defence, cutin development, and structural development of cell membranes and cell walls (Salminen *et al*., 2016). Both marker genes are highly relevant to the role of the pericarp as the main structural and protective layer of the seed.

The crease is a region of the wheat seed that carries a high bacterial load (Robinson *et al*., 2016), therefore pathogen defence and immune response genes are important to this tissue area. The strongest marker gene identified for the crease clusters in our gene expression matrix was LOC123104148 encoding induced stolen tip protein (TUB8). Induced stolen tip proteins are not yet characterised in wheat but have been reported in other plant species. A recent study in tomato identified an induced stolen tip protein as being a potential effect triggered immunity protein (Yu *et al*., 2021), and its expression in the crease could indicate that this gene could be involved in immune response and defence against pathogens in the wheat seed.

The embryo is a unique tissue in the seed with a distinct gene expression profile. The strongest marker gene identified for the embryo was LOC123113644, encoding cupincin protein. Cupincin also hasn’t been studied in wheat, however, in rice cupincin has been reported as a member of the cupin protein family and as having metallo-protease activity, influencing seed vigour (Sreedhar & Tiku, 2016). Therefore, this gene could also represent a potential target for optimising wheat seed performance.

The ‘inner endosperm ring’ cluster consists of transfer cells encircling the crease, whose role is the transfer of proteins, starch, lipids and nutrients from the crease to the rest of the seed during development (Lopato *et al*., 2014). The strongest marker genes for this cluster were non-specific lipid transfer protein genes and their high expression in this tissue during the peak of grain filling aligns with the anticipated expression patterns (Lopato *et al*., 2014). As these lipid transfer protein genes were some of the most highly expressed genes in the data overall, it is evident that they play a significant role in mid seed development and could be beneficial target genes for enhancing wheat yield. This represents an important avenue for further research.

The starchy endosperm cells of the sub-aleruone, central endosperm and inner endosperm boundary clusters make up the main part of the seed that is consumed by people, therefore genes highly expressed in these tissues could be relevant to the nutritional and baking quality of wheat. The strongest marker genes identified for both the sub-aleurone and central endosperm clusters were genes for α-, β- and γ-gliadins, storage proteins common to the endosperm and that play a key role in gluten network formation (Chaudhary *et al*., 2022). These proteins are an important source of nitrogen and amino acids and, along with glutenins, impact the texture and quality of bread dough (Barak *et al*., 2014). Similarly, the genes for α-amylase/trypsin inhibitor proteins were identified as marker genes for the inner endosperm boundary cluster. As α-amylase is an enzyme responsible for starch degradation (Call *et al.*, 2021), high expression of α-amylase inhibitors prevents this breakdown and ensures that starch accumulates sufficiently during grain filling (Ma *et al*., 2023). Due to their contribution to the functional and nutritional properties of wheat, these genes are already well characterised in the literature, however, their importance as endosperm specific genes that are highly expressed during the peak of grain filling is further highlighted by our study.

Many of the marker genes identified represent important targets to better understand the tissue-specific processes and interactions taking place during mid seed development. As different tissues and cellular groups contribute to different developmental traits, detail about the most highly expressed genes in those regions will allow for more targeted genetic research on those traits.

## Conclusion

This study confirmed the spatial expression patterns of known tissue specific genes and identified novel marker genes for the pericarp, crease, embryo and endosperm tissues of the developing wheat grain. These findings also highlighted the utility of spatial transcriptomics in studying subgenome differences, as we were able to distinguish subgenome biased expression, and subgenome specific spatial expression patterns in a polyploid plant. As different tissues and cellular groups contribute differently to the functional and nutritional properties of seeds, spatial detail of gene expression during development provides valuable information for fine tuning targeted genetic research. The nutritional importance of seeds, particularly cereal grains, means that spatial gene expression data marks a significant advancement in our capacity to optimise food resources.

## Material and Methods

### Plant growth conditions and sample collection

Bobwhite SH98 26 seeds were germinated and grown in pots, 5 seeds per pot, in a glasshouse during winter in Brisbane, Australia. They were grown in media containing 2g/L of Osmocote 3-4 month fertiliser and watered twice daily for 2 min at a time. Plants were monitored for the emergence of the first spikelet at 2.5 months of growth. Once the beginning of spikelet emergence could be observed, plants were tagged and dated, allowing for seed collection at 14DPA, the stage of development where grain filling is at its peak (Kim *et al*., 2024).

Seeds were collected from the central third of the wheat ears, the glumes and lemmas were removed, and the seeds immediately transferred into ambient temperature (liquid) Optimum Temperature Compound (OCT), in 1x1 cm containers. All seeds were placed on the bottom of the containers with the crease facing downwards, so they remained on the same plane. The containers were then transferred straight to dry ice where the OCT quickly froze. Sterile forceps were used to arrange the seeds and prevent them from floating up in the OCT before the block fully set. The time from removal of seeds from the plant to freezing was under 20 s.

### Sample preparation and tissue mounting

Seed sections were cut using a cryostat (CryoStar Nx70, Epredia). The cryostat chamber was cleaned with 70% ethanol and cooled to -22°C, and the blade to -18°C. The samples were placed in the chamber and allowed to equilibrate to -22°C for 30 min, as well as forceps and paint brushes for handling the tissue.

Stereo-seq Capture Chips were removed from vacuum sealed bags and allowed to equilibrate to room temperature. To prepare the chips for mounting, they were rinsed with 100µl of nuclease free water and then dried by power duster. Then 50µl of 0.01% poly-l-lysine was added dropwise to the chip surface and left to stand for 10 min. The poly-l-lysine was then removed by power duster and the chips again rinsed with 100µl of nuclease free water and dried. Chips were then placed in the cryostat chamber and cooled to -22°C.

Once cooled for 30 min, the samples were mounted onto the specimen disk holder and trimmed to the section to be analysed. Here, that was approximately one third of the way through the seeds when cutting longitudinally, where both the crease and embryo were included in the section. Tissue sections were cut at 20µm thick and transferred to Stereo-seq Capture chips using chilled forceps and brushes.

To mount the tissue sections, the chips were picked up within the cryostat chamber and warmed by pressing a finger to the back of the slide. The OCT thawed and the tissue section adhered to the chip surface. The slide was then dried at 37°C for 1 min and fixed in methanol at -20°C for 30 min.

### Permeabilization experiment

The permeabilization experiment was conducted first to determine the ideal permeabilization time for this tissue type, which would later be used for the transcriptomics experiment. This was done using the Stereo-seq Permeabilization Set User Manual and following the manufacturer’s guidelines with little modifications (BGI, 211SC114) (Chen *et al*., 2022; Xia *et al*., 2022). In brief, four seed samples were each mounted onto a Stereo-seq P chip and fixed (as described previously). After fixation the samples were incubated at 37°C in 0.1% PR enzyme in 0.01M HCl buffer, for 6, 12, 18 and 24 min respectfully. The samples were then washed with 0.1x saline sodium citrate (SSC) solution. Reverse transcription was performed for 1 hr at 42°C using the RT QC Mix, and then the tissue was removed from the chip surface by Tissue Removal Buffer.

Fluorescent images were taken of the chips at 5x and 10x objective lenses, in the red channel. Based on the images taken (Figure S22), 12 min produced the brightest image with the least degradation, and was determined to be the optimal permeabilization time for wheat grain tissue.

### Transcriptomics experiment

The transcriptomics experiment was conducted using the Stereo-seq Transcriptomics Set User Manual and following the manufacturer’s instructions with little modification (BGI, 211SC114) (Chen *et al*., 2022; Xia *et al*., 2022). In brief, four seed samples were each mounted onto a Stereo-seq T chip and fixed (as described previously). After tissue fixation, the sections were stained for fluorescent imaging by the Qubit ssDNA reagent. Fluorescent images were taken of each chip using the 10x objective lens in both the FITC channel and DAPI channel.

The samples were then permeabilised in 0.1% PR enzyme in 0.01M HCl buffer for 12 minutes at 37°C, so that the RNA could be released from the tissue sections and captured on the chips’ surface. Reverse transcription was then performed overnight at 37°C. The next day, Tissue Removal Buffer was used to remove the tissue from the chips’ surface and the remaining cDNA was released by cDNA Release Buffer for 3 hours.

The collected cDNA was then used for library construction by the Stereo-seq Library Preparation kit. The resulting libraries were pooled and sequenced by whole genome sequencing using the T7 PE100 flow cell.

During the protocol, challenges arose with the tissue mounting step, resulting in some folding and breaking of the tissue sections. We proceeded with the most intact tissue sections for the subsequent analyses.

### SAW Pipeline

The SAW software suite (https://github.com/BGIResearch/SAW), with default parameters, was used to map the raw sequencing reads to the wheat reference genome IWGSC CS RefSeq v2.1 (Zhu *et al*., 2021). Reads that could be uniquely mapped to coding regions of the genome were counted, and the coordinate identity (CID) of each read was used to place it spatially in its original position on the chips. This data was overlayed with the fluorescent microscope images of the tissue sections before permeabilization, so that reads could be mapped to their spatial location within the tissue. The SAW software suite was then used to generate a matrix of expression count of all genes expressed at each spot on the chips. To handle the sparseness of the gene expression within each spot (i.e. only a subset of genes are captured per spot) adjacent spots could be grouped into different bin levels, such as Bin200 for 200x200 spots, for analysis at varying resolutions. It was determined that Bin50 was the ideal resolution for this data.

Once the gene expression matrices were generated, one tissue section was deemed to be the highest quality replicate compared to the others which exhibited damage due to challenges during the tissue mounting step. The chip containing the most intact replicate was designated as the focus for the subsequent analysis, while the remaining chips were referenced to provide replication where possible.

### Clustering and marker gene discovery

The Stereopy library was used to generate gene clusters and marker genes at the desired Bin level, Bin50. The gene expression matrices were first filtered to remove bins with low MID counts. Due to the expression levels of genes within the tissue boundaries and expression levels of the background noise outside the tissue boundaries, a threshold of minimum 30 MID counts per bin was selected for the focus chip, and a threshold of between minimum 30 and 50 MID counts was selected for the replicate chips. Filtering the data by expression levels also increased the computational efficiency of the analysis and allowed for downstream generation of spatial neighbours clusters.

The expression data was normalised by the log1p method (Booeshaghi & Pachter, 2021) and then underwent principal component analysis (PCA). A neighbourhood graph of bins was then generated using the PCA representation of the expression matrix, followed by a spatial neighbours graph. These graphs were then embedded into two dimensions for ease of visualisation using UMAP.

Leiden clustering was then undertaken using the spatial neighbours data, and 11-12 distinct gene expression clusters were identified for each chip. Overlaying the spatial Leiden clusters with the fluorescent microscope images of the tissue allowed us to link eight clusters to functional cellular groups and determine that some clusters could be discarded as background noise or as a result of tissue folding issues. This was due to their location outside the tissue boundary, or a pattern following the shape of the folded areas that did not align with any particular cell or tissue type.

Lastly, marker gene candidates were identified for each gene expression cluster, using a t-test on the raw MID count data, to compute a ranking of differentially expressed genes across clusters. This analysis was repeated for all chips analysed, and marker genes identified for their tissue type on multiple chips were recorded.

### Investigation of known tissue-specific genes

Known tissue specific genes were identified from the literature, including genes expected or known to be specifically expressed in the endosperm, pericarp and embryo tissues, and Blastn analysis was used to identify subgenome paralogs (Sayers *et al*., 2025). Their spatial expression patterns and subgenome specific expression were investigated using our gene expression matrix and the Stereomap app. The expression of homologous gene triplicates were visualised in groups and individually, in order to capture their overall expression and compare between subgenomes.

Furthermore, a sub-sample of 5% of bins from each cluster were randomly selected and used to undertake differential expression analyses. The bins were selected from clusters generated using the log1p normalised expression data, and those bin IDs were then used to extract the unnormalized MID count data from those locations in the chip. The raw count data for the tissue specific target genes at each randomly sampled bin ID (location) were then used for differential expression analysis to quantify the differences in expression of homologous gene copies across different clusters (tissue types) and also to quantify the differences in overall expression between pairs of genes within homologous gene triplicates. Lastly, we also compared expression of individual α-amylase/subtilisin inhibitor paralogs across clusters, as these genes appeared to exhibit spatial subgenome biased expression, based on the Stereomap images (Figure 4DEF).

To determine the statistical significance of the differences in gene expression we used an approximate (Monte Carlo) permutation test. This test was chosen as the highly zero-inflated spatial expression data would violate the normality assumption of a standard t-test and produce poor results under an equivalent rank-based test. In the approximate permutation test, 500,000 random permuted samples of the two sets of gene expression data being tested were generated, which provided an estimate of the p-value. Binomial theory was then applied to give a 99.9 % confidence interval for the true p-value. The estimated p-values were then adjusted by a Bonferroni multiple testing correction factor of 264, the total number of tests conducted, resulting in values that we have a high level of confidence the true adjusted p-values were less than, as reported in Tables S1-S9.

## Acknowledgements

This project was supported by the ARC Centre of Excellence for Plant Success in Nature and Agriculture (CE 200100015).

## Author contributions

RH conceived and supervised the project. TM, RS, KXS and KLL conducted the experiments. TM, DK, LM and AM conducted the analysis. TM wrote the manuscript with input from DK, RS and RH. All authors read the final version of the paper.

## Data availability

The high-throughput data that supports the findings of this study are openly available in Gene Expression Omnibus at https://www.ncbi.nlm.nih.gov/geo/query/acc.cgi?acc=GSE298021, accession number GSE298021. The scripts used have been uploaded to GitHub (https://github.com/tori-millsteed/Wheat_seed_spatial_transcriptomics_Millsteed2025).

## Supporting information

Supporting data file 1: *supporting_data_1_additional_figures.docx*

Supporting data file 2: *supporting_data_2_permutation_tests.xlsx*

