## Supporting data file 1 for "Spatial transcriptomics of developing wheat seed reveals radial expression patterns in endosperm and subgenome biased expression of key genes"

### Key Metrices

| Metric | Reads | % |
|---|---|---|
| Total Reads | 780,145,171 | 100% |
| Valid CID Reads | 570,305,187 | 73.1% |
| Clean Reads | 542,585,536 | 95.1% |
| Uniquely Mapped Reads | 36,415,138 | 6.7% |
| Transcriptome | 16,611,992 | 45.6% |
| Unique Reads | 5,741,511 | 34.6% |
| Sequencing Saturation | 10,870,481 | 65.4% |
| Unannotated Reads | 19,803,146 | 54.4% |
| Multi-Mapped Reads | 493,843,278 | 91.0% |
| Unmapped Reads | 12,327,120 | 2.3% |
| Non-Relevant Short Reads | 27,719,651 | 4.9% |
| Invalid CID Reads | 204,300,199 | 26.2% |
| Discarded MID Reads | 5,539,785 | 0.7% |

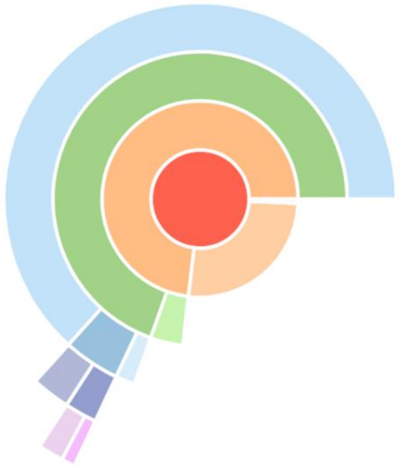

**Figure S1.** SAW analytical report key metrices for chip 1 (chip ID D02266B1)

### Tissue

| | |
|---|---|
| DNB Under Tissue | 92,046,597 |
| mRNA-Captured DNBs Under Tissue | 3,273,120 |
| Genes Under Tissue | 60,877 |
| Number of MID Under Tissue Coverage | 4,244,343 |
| Fraction MID in Spots Under Tissue | 73.92% |
| Reads Under Tissue | 365,103,798 |
| Fraction Reads in Spots Under Tissue | 63.40% |

**Figure S2.** SAW analytical report tissue data for chip 1 (chip ID D02266B1)

### Key Metrices

| | | |
|---|---|---|
| Total Reads | 1,110,260,180 | 100% |
| Valid CID Reads | 857,718,889 | 77.3% |
| Clean Reads | 824,477,468 | 96.1% |
| Uniquely Mapped Reads | 79,536,921 | 9.6% |
| Transcriptome | 52,673,637 | 66.2% |
| Unique Reads | 14,320,878 | 27.2% |
| Sequencing Saturation | 38,352,759 | 72.8% |
| Unannotated Reads | 26,863,284 | 33.8% |
| Multi-Mapped Reads | 734,027,333 | 89.0% |
| Unmapped Reads | 10,913,214 | 1.3% |
| Non-Relevant Short Reads | 33,241,421 | 3.9% |
| Invalid CID Reads | 244,232,972 | 22.0% |
| Discarded MID Reads | 8,308,319 | 0.7% |

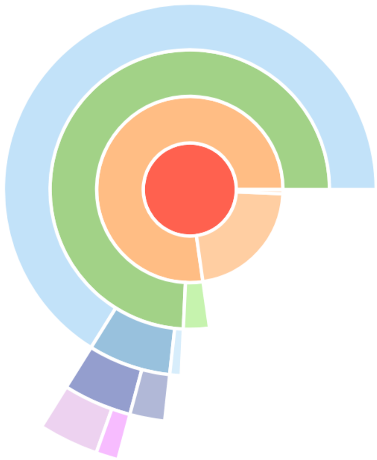

**Figure S3.** SAW analytical report key metrices for chip 2 (chip ID D02266A4)

### Tissue

| | |
|---|---|
| DNB Under Tissue | 90,326,917 |
| mRNA-Captured DNBs Under Tissue | 7,382,161 |
| Genes Under Tissue | 60,916 |
| Number of MID Under Tissue Coverage | 11,599,308 |
| Fraction MID in Spots Under Tissue | 81.00% |
| Reads Under Tissue | 507,757,235 |
| Fraction Reads in Spots Under Tissue | 58.63% |

**Figure S4.** SAW analytical report tissue data for chip 2 (chip ID D02266A4)

### Key Metrices

| | | |
|---|---|---|
| Total Reads | 1,324,226,135 | 100% |
| Valid CID Reads | 1,010,302,264 | 76.3% |
| Clean Reads | 973,284,122 | 96.3% |
| Uniquely Mapped Reads | 49,272,216 | 5.1% |
| Transcriptome | 22,842,830 | 46.4% |
| Unique Reads | 6,036,443 | 26.4% |
| Sequencing Saturation | 16,806,387 | 73.6% |
| Unannotated Reads | 26,429,386 | 53.6% |
| Multi-Mapped Reads | 911,428,823 | 93.6% |
| Unmapped Reads | 12,583,083 | 1.3% |
| Non-Relevant Short Reads | 37,018,142 | 3.7% |
| Invalid CID Reads | 305,429,088 | 23.1% |
| Discarded MID Reads | 8,494,783 | 0.6% |

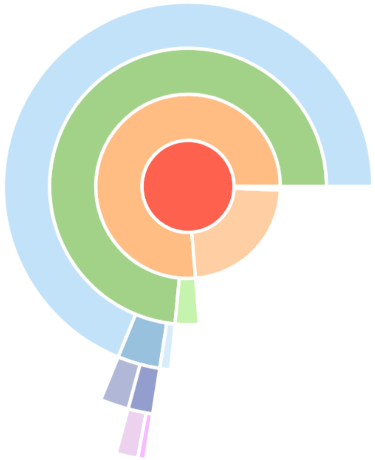

**Figure S5.** SAW analytical report key metrices for chip 3 (chip ID D02263D4)

### Tissue

| | |
|---|---|
| DNB Under Tissue | 85,660,333 |
| mRNA-Captured DNBs Under Tissue | 3,625,784 |
| Genes Under Tissue | 59,190 |
| Number of MID Under Tissue Coverage | 5,218,664 |
| Fraction MID in Spots Under Tissue | 86.45% |
| Reads Under Tissue | 574,757,616 |
| Fraction Reads in Spots Under Tissue | 56.42% |

**Figure S6.** SAW analytical report tissue data for chip 3 (chip ID D02263D4)

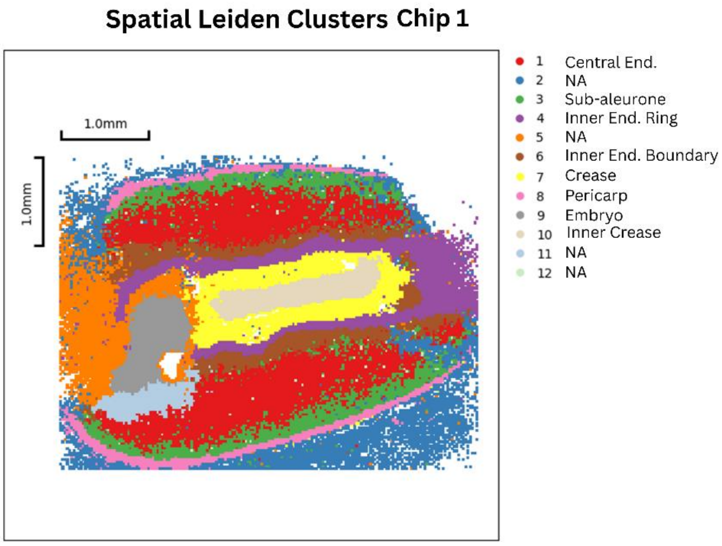


**Figure S7:** Spatial Leiden clusters for chip 1 (zoomed in to view only one seed section)

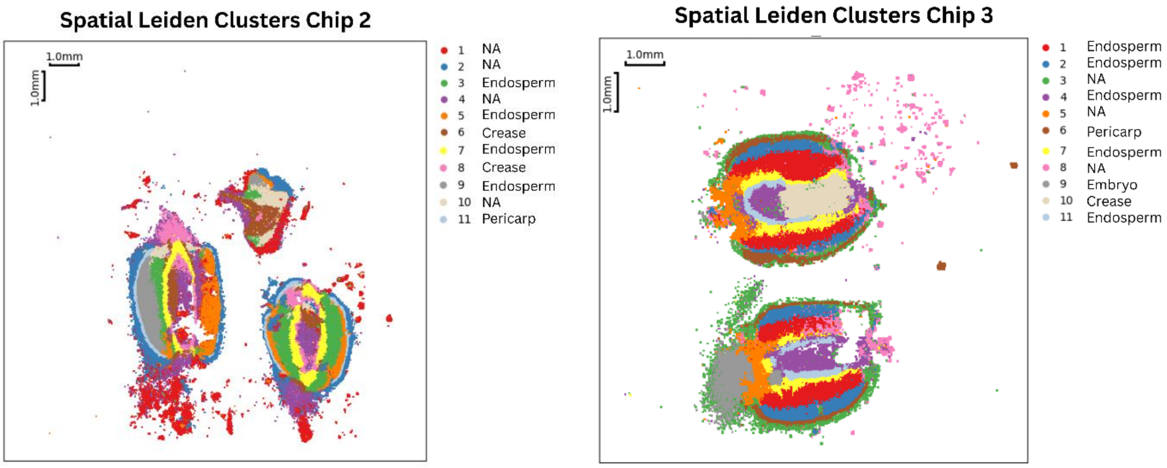


**Figure S8:** Spatial Leiden clusters of chips 2 and 3, no zoom applied

TaNAC019

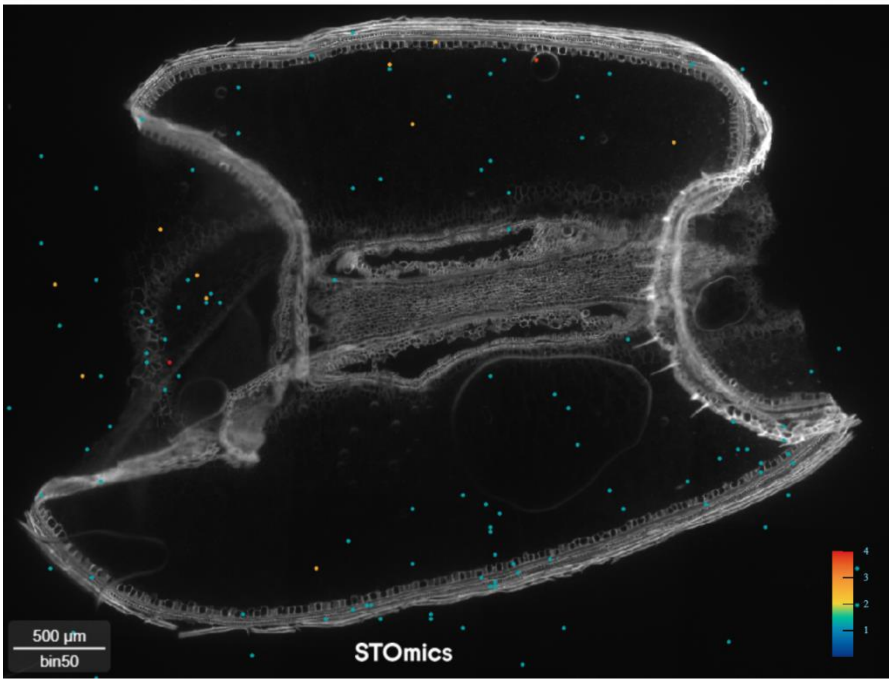


**Figure S9.** TaNAC019 spatial expression

TabZIP28

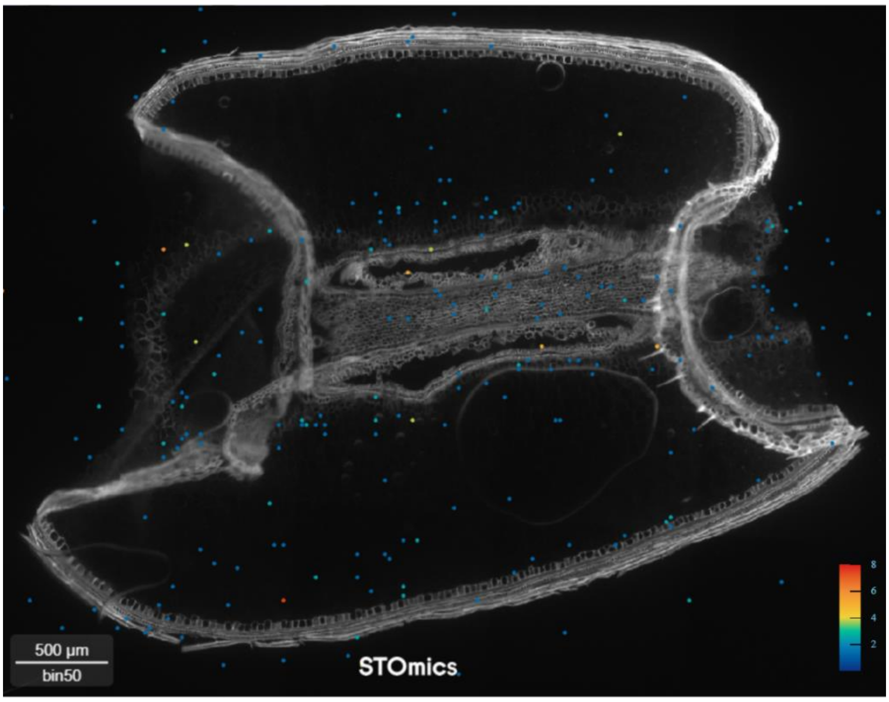


**Figure S10.** TabZIP28 spatial expression

Metallothionein-protein

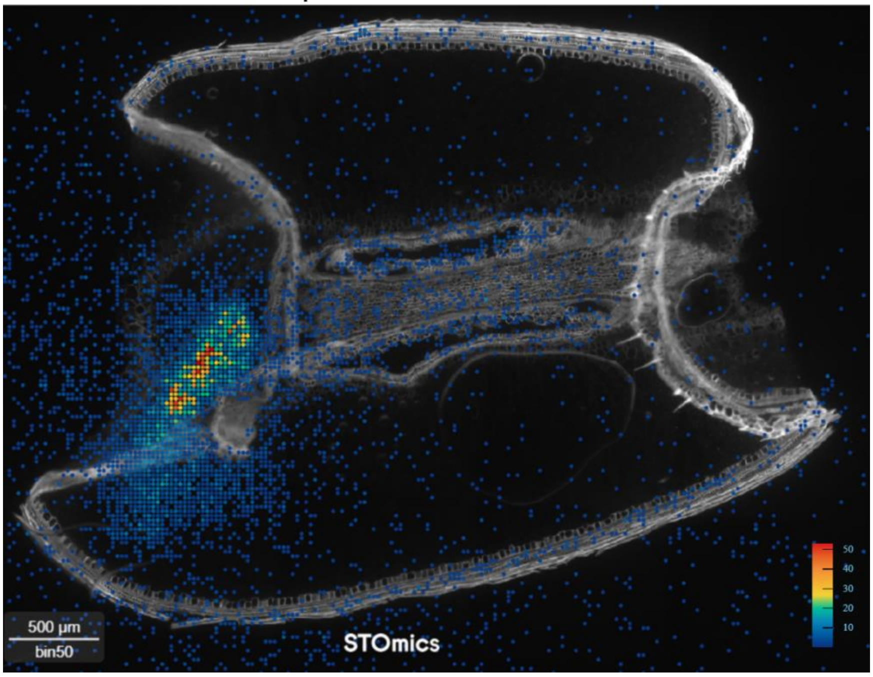


**Figure S11.** Metallothionein-protein spatial expression

EM Promoter protein

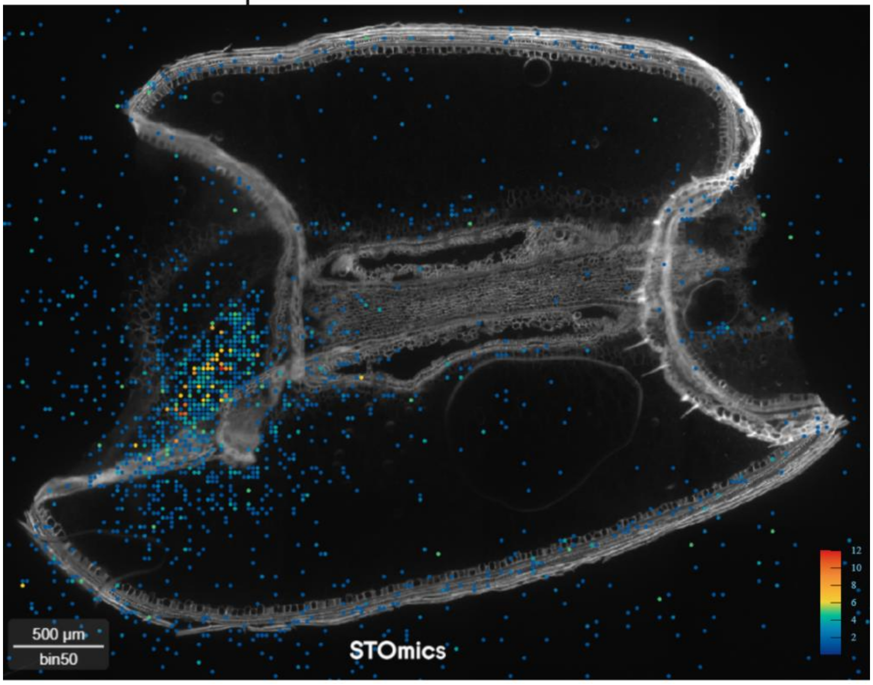


**Figure S12.** EM promoter protein spatial expression

Pyruvate orthophosphate dikinase

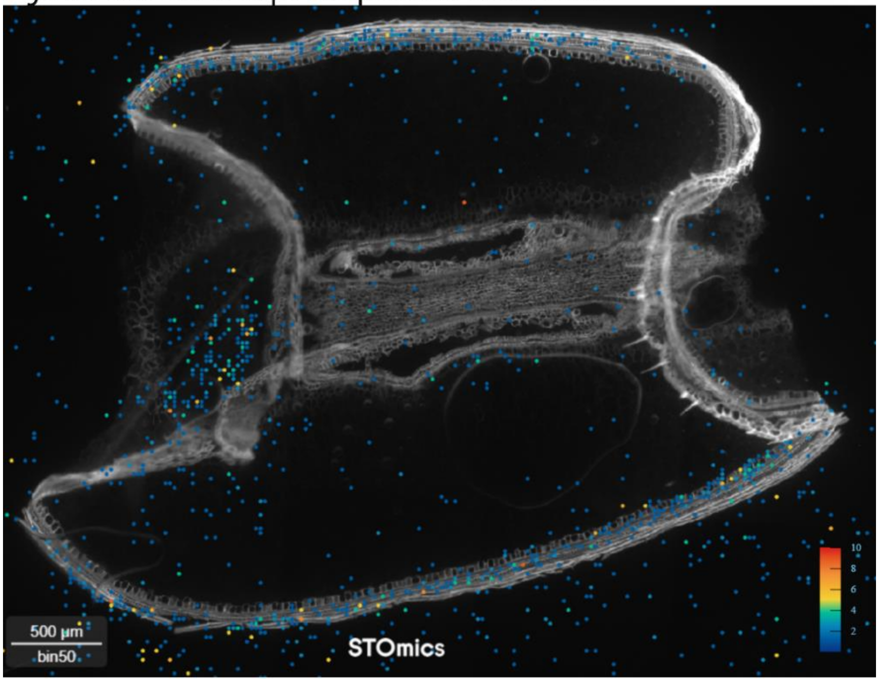


**Figure S13.** Pyruvate orthophosphate dikinase spatial expression

Subgenome A Subgenome B Subgenome D

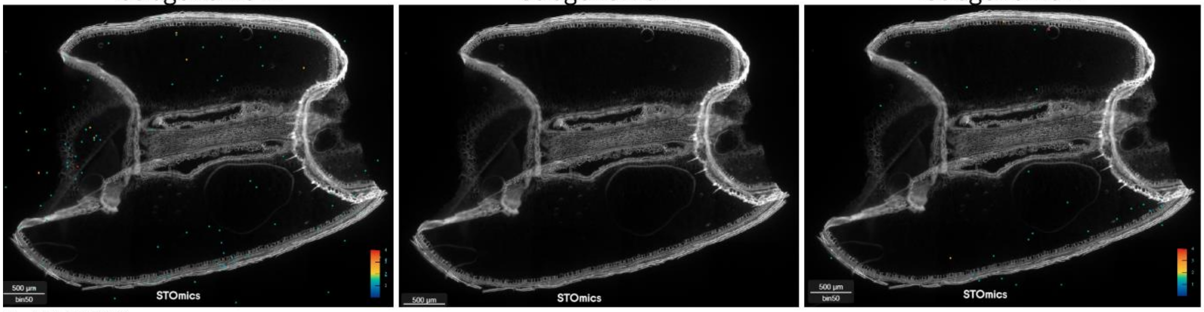


TaNAC019

**Figure S14.** TaNAC019 subgenome specific expression

Subgenome A Subgenome B Subgenome D

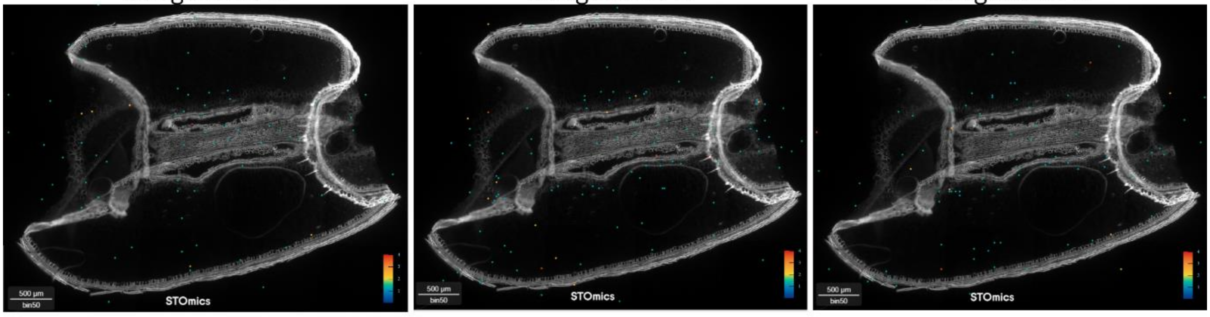


TabZIP28

**Figure S15.** TabZIP28 subgenome specific expression

Subgenome A Subgenome B Subgenome D

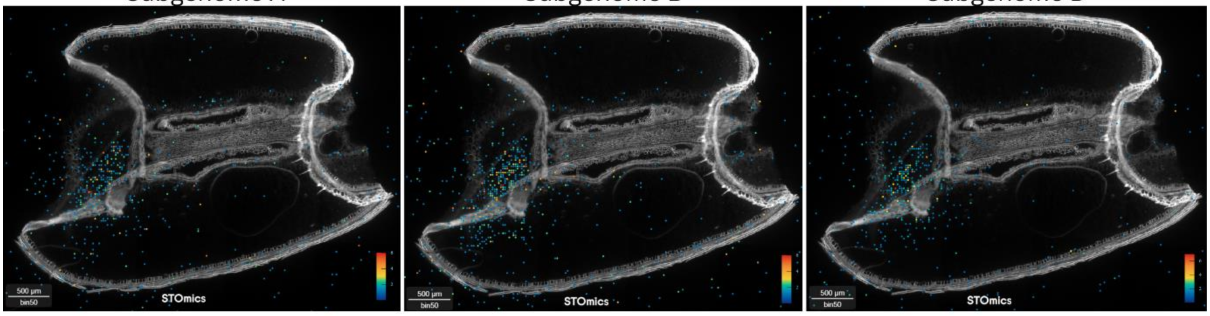


EM promoter protein

**Figure S16.** EM promoter protein subgenome specific expression

Subgenome A Subgenome B Subgenome D

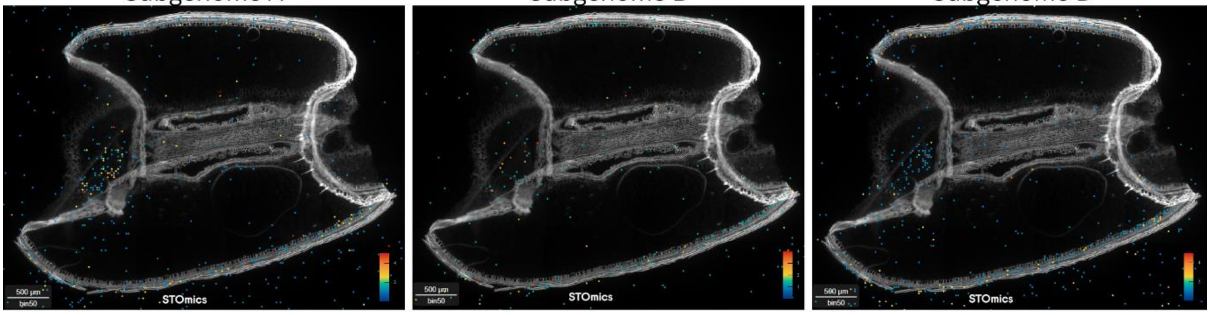


Pyruvate orthophosphate dikinase

**Figure S17.** Pyruvate orthophosphate dikinase subgenome specific expression

Subgenome A Subgenome B Subgenome D

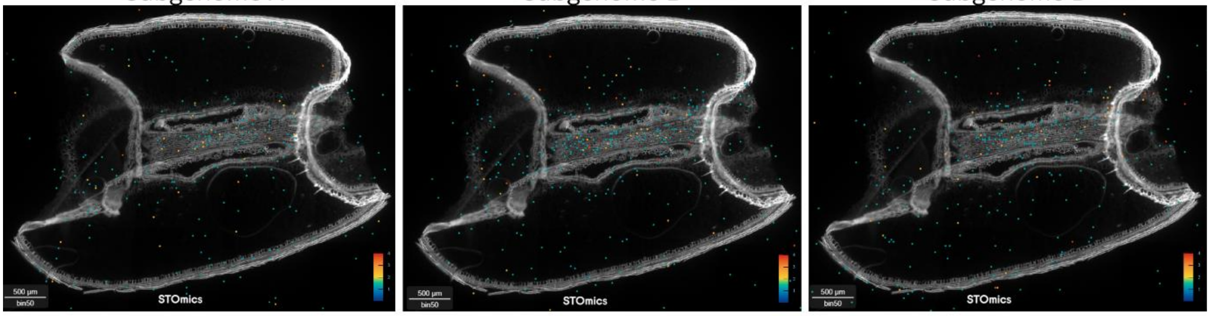


Autophagy related protein 8ATG

**Figure S18.** Autophagy related protein 8ATG subgenome specific expression

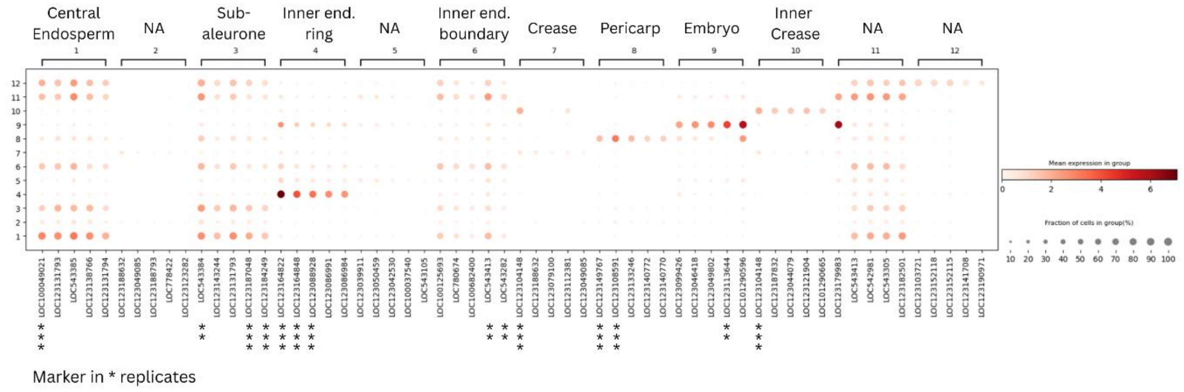


**Figure S19.** Marker genes plot of chip 1

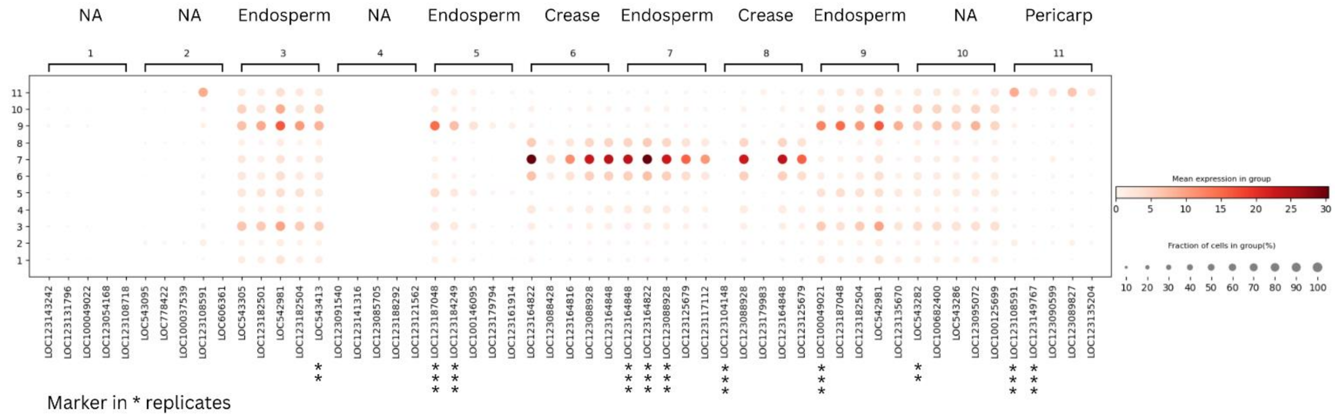


**Figure S20:** Marker genes plot of chip 2

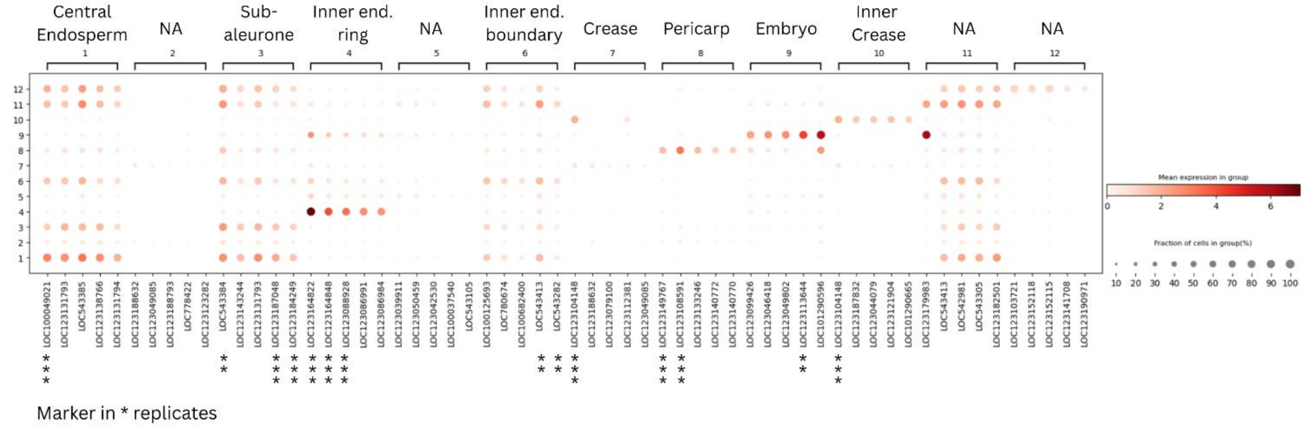


**Figure S21:** Marker genes plot of chip 3

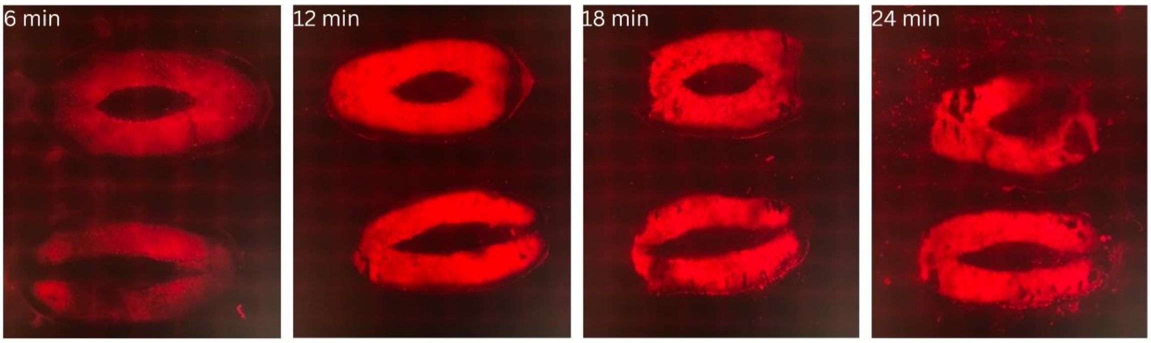


**Figure S22.** Permeabilization results after 6, 12, 18 and 24 min of treatment with permeabilization reagent
